# The Effects of CdSe/ZnS Quantum Dots on the Photosynthesis Rate of the *Chlorella Vulgaris* Beads

**DOI:** 10.1101/2022.08.01.500842

**Authors:** Wimeth Dissanayake, Richard Hailstone, Mengdi Bao, Ruo-Qian Wang, Xin Yong, Ke Du

## Abstract

Photosynthesizing microalgae produce more than 50% of oxygen in the atmosphere and are crucial for the survival of many living systems such as coral reefs. To address the declining of coral reefs, artificial reefs have been introduced to encapsulate the aglae cells in a polymer matrix but the effects of nanoscale pollutants on these engineered systems have not been fully understood. In this work, quantum dots with a size smaller than 10 nm are being used to elucidate the photosynthesis performance of the sodium alginate beads encapsulated with *Chlorella vulgaris* (*C. vulgaris)*. The fluorescent quantum dots can move into the alginate matrix and the fluorescence intensity in the algae beads is correlated with the quantum dot concentration. We further show that the photosynthesis of the algae beads are sensitive to the quantum dot concentration and are also time sensitive. In the first 48 min of quantum dot exposure, both carbon dioxide absorption and oxygen production are low, suggesting limited photosynthesis. After the initial incubation, the photosynthesis rate quickly increases even though more inhibition is still observed with higher concentration of the quantum dots.

## Introduction

Green algae, such as *Chlorella vulgaris* (*C. vulgaris*), has shown great potential to be used in pollution control^1^, biofuel^2^, and dietary supplement^3^. Containing chloroplasts, algae can photosynthesize and produce half of the atmospheric oxygen^4^. They are also crucial to the marine systems via the formation of symbiotic relationships with other organisms, particularly coral reefs. Coral reefs play an essential role in shoreline protection by reducing the wave energy by 97%^5^. However, the reefs have decreased by ∼50% since 1950 due to rapid global warming, thus increasing the risks of global natural hazards^6^. A recent study has shown that heat-evolved algae can endure elevated temperatures and enhance coral bleaching tolerance to marine heat waves^7^. To further enhance the photosynthetic efficiency of coral reefs and potentially create artificial living reefs to combat climate change, 3-D printed structures containing algae cells have been introduced to mimic the morphological features of coral reefs^8,9^. Alginate hydrogel shows great promise as the scaffold material, in which the cells are viable for several days without nutrition^10^.

Another emerging challenge for the coral reefs and future engineering living reefs is the continuous exposure to chemical waste, especially at nanoscale^11,12^. Early results reported decreases in cell growth rate and chlorophyll content when the cell cultures were exposed to nanomaterials such as quantum dots^13^, nanoplastics^14^, and oxide nanoparticles^15^. On the other hand, gold nanoparticles can enhance photosynthesis in microalgae by transferring light into photogenerated electrons. A 42.7% increase in the carotenoids has been reported with this method^16^. To date, most of the research on nanomaterials/microalgae interactions has focused on cell cultures in solution, and little has been investigated with the immobilized microalgae cells that can be constructed as engineering living systems.

Here, we use quantum dots as a model to study the interaction of nanostructures with engineering living systems by encapsulating *C. vulgaris* in sodium alginate beads (**Figure 1a**). We show that more quantum dots are presented in the algae beads as the quantum dot concentration in the solution increases. We apply the bicarbonate indicator and gas chromatography-mass spectrometry to study the effect of quantum dots on algae photosynthesis. In the first 48 min after incubation, the photosynthesis is significantly inhibited by the presence of quantum dots. However, the photosynthesis rate increases after the initial incubation, suggesting that some of the quantum dots might be released or digested by the algae cells. At 120 min, algae beads incubated with quantum dots show less oxygen production than the samples without quantum dots, and the oxygen production is inversely correlated with the quantum dot concentration. The results indicate that the microalgae-based living systems are sensitive to the environment with nanoscale pollutants.

**Figure 1.**
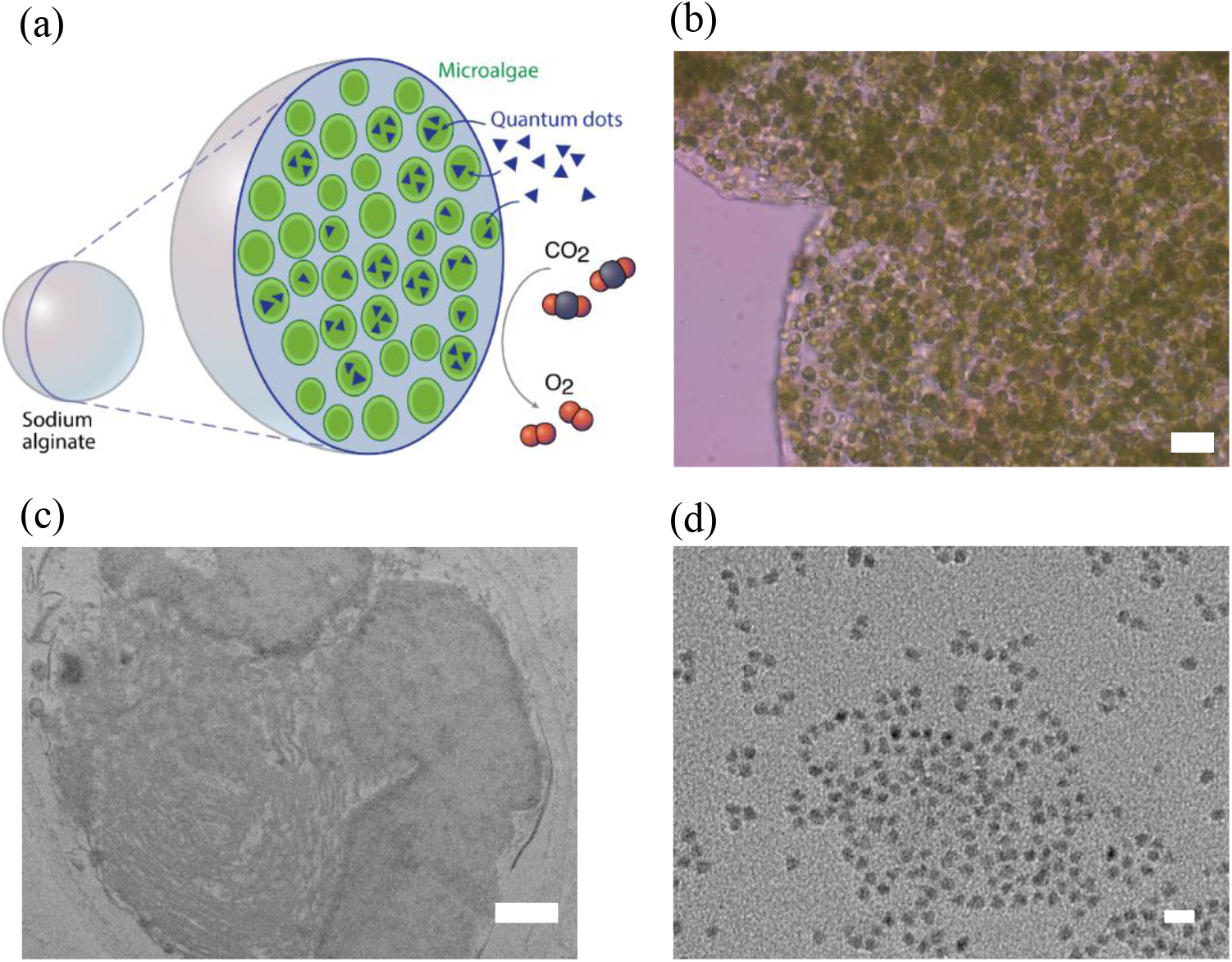
(a) The schematic showing the endocytosis and photosynthesis process of microalgae beads with quantum dots. (b) Bright-field image of the algae beads (scale bar: 20 µm). TEM image of (c) Microalgae cell (scale bar: 400 nm) and (d) Quantum dots (scale bar: 10 nm).

## Materials and Methods

### Materials

*C. vulgaris* beads were obtained from Algal Research Supply and were manufactured using Sodium Alginate and Calcium Chloride^17^. The bright field and TEM image of a disseted bead are shown in **Figure 1b** and **1c**, respectively. Algal cultures were purchased from Algal Research Supply and grown using Bold’s Basal Medium. Streptavidin - ZnS/CdSe quantum dots (5 nm size), with 525 nm emission maxima, were purchased from Thermofisher Scientific (**Figure 1d**). Philips T12, 40-watt, cool white fluorescent lights were used to incubate and grow the microalgae cells on a 12hr/12hr light-dark cycle.

### Incubation of microalgae beads with quantum dots

Ten microalgae beads were placed into a 200 µL PCR tube with 100 µL of 0 µM, 0.01 µM, 0.05 µM, and 0.1 M of quantum dot solution. The solution was first mixed for 2 min, then centrifuged at 1500 rpm for 15 min, followed by 90 min of room temperature incubation under a T12, cool white, fluorescent light.

### Bright-field and fluorescence imaging

The microalgae bead was dissected into hemispheres and prepared on a wet glass slide with a cover. The slide was then viewed using the Bright Field mode of an Amscope XD-RFL microscope. Bright-field imaging was used to identify clusters of the *C. vulgaris* cells and to quantify the relative concentration of the cells. The Amscope XD-RFL was also used to image the slides fluorescently.

### TEM imaging

Intact microalgae beads were first fixed with 2% glutaraldehyde in 0.2 M cacodylate buffer for 2 hr. They were then rinsed three times (15 min each) in 0.2 M cacodylate buffer. They were then postfixed in 1% OsO_4_ in 0.2 M cacodylate buffer for two hours, followed by three rinses (15 min each) in 0.2 M cacodylate buffer. The beads were then dehydrated in a series of increasing concentrations (25%, 50%, 75%, and 95%) of ethanol for 10 min at each concentration. Finally, the beads were immersed in 100% acetone for 10 min. The dehydrated beads were immersed in 3/1 acetone/Embed 812 (Electron Microscopy Sciences) for 12 hr, followed by 1/1 acetone/Embed 812 for four hr, followed by 3/1 acetone/Embed 812 for four hr. The beads were then embedded in Embed 812 for 12 hr, followed by polymerization at 60 ^°^C for 12 hr.

Thin sections (70 nm) for imaging were made with a Leica Artos 3D ultramicrotome. The thin sections were floated on water and transferred to 150 mesh copper grids. Staining was performed with UranyLess EM stain (Electron Microscopy Sciences) for two min, followed by two 15 s rinses in distilled water. Imaging was performed with a JEOL 2010 transmission electron microscope operating at 200 kV. Images were captured with an AMT digital camera.

### Quantification of fluorescence

The fluorescence images were quantified using Fiji ImageJ. Several areas of cells containing quantum dot fluorescence were analyzed to find integrated density. Several areas of background were analyzed to find mean background fluorescence. Thereby the ImageJ program could calculate a baseline for fluorescence which could be used to correct the average fluorescence found from the selected areas of note, using the equation:

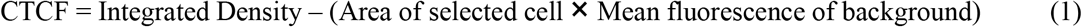

The area of fluorescence was identified by using threshold values of 83 and 87 HSG, the values most closely matched the emission of the quantum dots, to isolate the fluorescence of the quantum dots. After isolation, the area of these regions was integrated to obtain a total area of fluorescence.

### Production of CO_2_

Stored CO_2_ was generated by the chemical reaction:

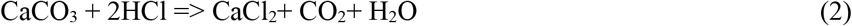

Two 4-liter sealable glass containers were connected by Teflon tubing. We added 22 mL of HCl to one of the containers and followed by another 47g of CaCO_3_. The glass was then promptly sealed, only allowing gas to pass through the Teflon tubing into the second container, separating CO_2_ from other products. Using a thin 100 µL Hamilton syringe, 3.6 L of CO_2_ gas was extracted from the second container for later use.

## Results and Discussions

The microalgae beads are large in size (diameter: 1.5 mm) and tend to sediment to the bottom of the microtube. We exploited a short and low-speed centrifuge process to facilitate the interaction between quantum dots and algae beads. As shown in **Figure 2**, CTCF values of 28,331 and 37,856 are achieved by two different centrifuge processes of 950 rpm for 3 min and 1,500 rpm for 5 min, respectively. While previous studies showed absorption of quantum dots in free-floating *Chlamydomonas* with only 960 rpm for 3 min^13^, fluorescence imaging of the dissected *Chlorella* beads showed minimal absorption into the cells with this protocol. Compared to *Chlamydomonas* species, *Chlorella* species have a much more resilient cell wall due to the “algaenan” outer coating of their cell walls^18^. Other studies reported that free-floating *Chlorella* absorbed high concentrations of quantum dots when centrifuged at 13,400 rpm for 15 min^19^. However, this high speed would damage the alginate substrate holding *Chlorella* cells in this study. Therefore, we selected the 1,500 rpm/5 min process for all the following experiments.

**Figure 2.**
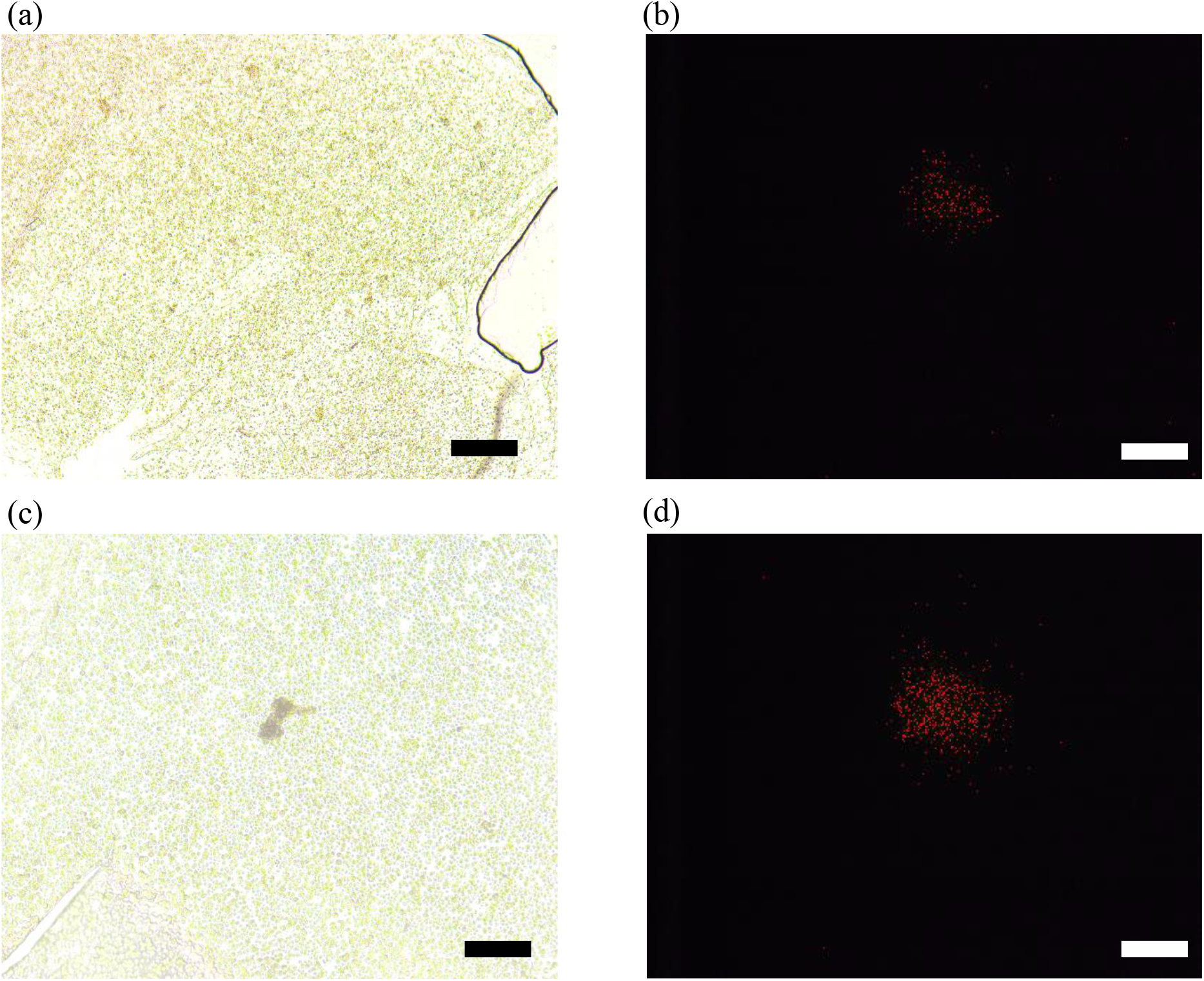
(a) Bright-field and (b) fluorescence image of *Chlorella vulgaris* cells encapsulated alginate beads centrifuged at 950 rpm for 3 min. The CTCF shown in (b) is 28,331. (c) Bright-field and (d) fluorescence image of the beads centrifuged at 1,500 rpm for 5 min. The CTCF shown in (d) is 37,856. The scale bar is 50 µm.

After establishing the incubation protocol, we then evaluated the fluorescence intensity of the microalgae beads versus quantum dot concentration (0, 0.01, 0.05, and 0.1 µM) by using both bright-field and fluorescence imaging. As shown in **Figure 3**, increasing the quantum dot concentration from 0 to 0.1 µM increases the CTCF value of the microalgae beads. The calculated CTCF value and the total fluorescence area shown in **Figures 4a** and **4b** both present a logarithmic relationship with the quantum dot concentration. This dependence indicates that the uptake capacity for the cells decreases with the increasing amount of the quantum dots.

**Figure 3.**
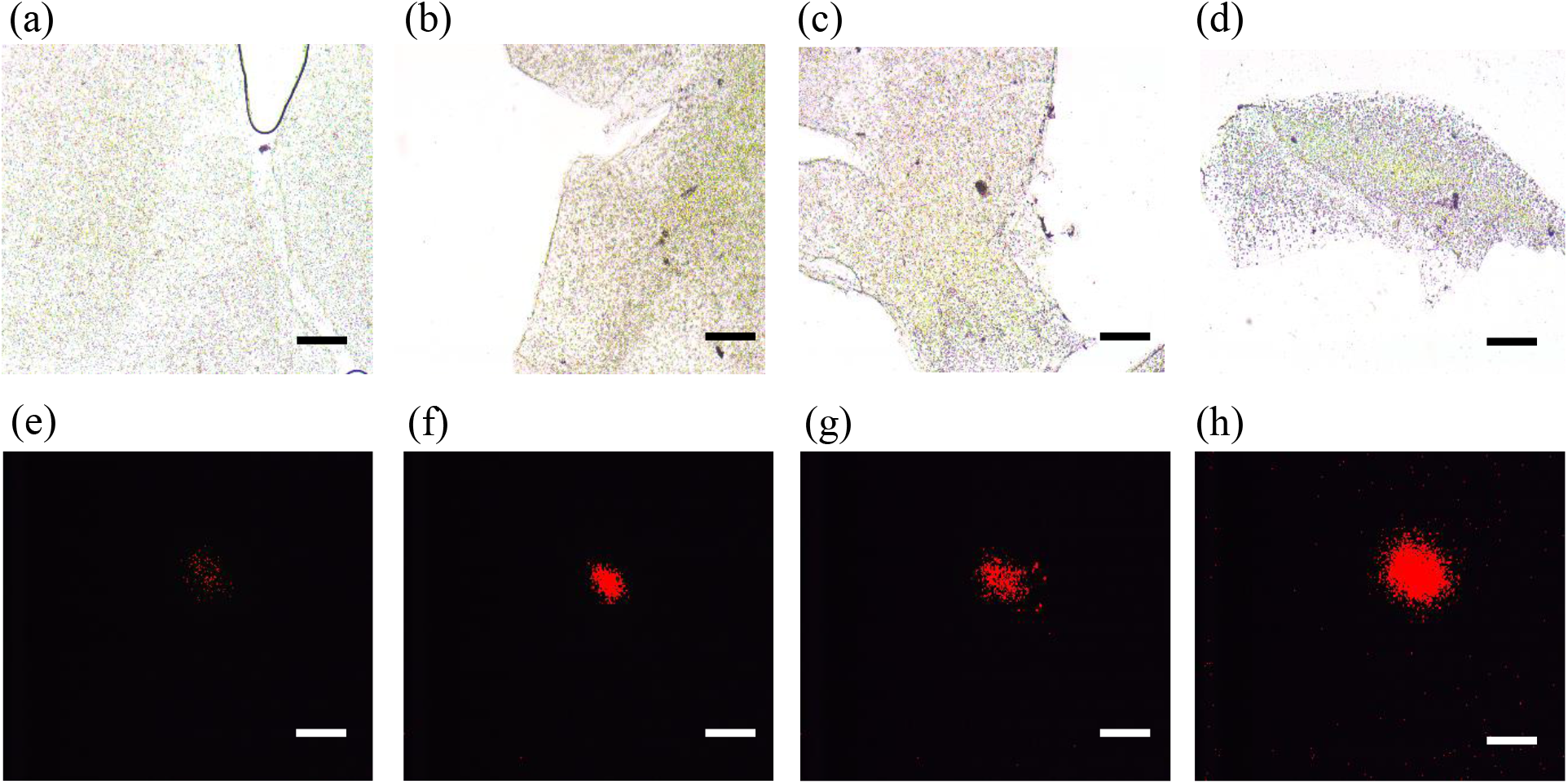
Bright-field and Fluorescent images of alginate beads incubated with different concentrations of quantum dots. (a) and (b): Control (0 µM). (c) and (d): 0.01 µM. (e) and (f): 0.05 µM concentration. (g) and (h): 0.1 µM. Scale bar is 50 µm.

**Figure 4.**
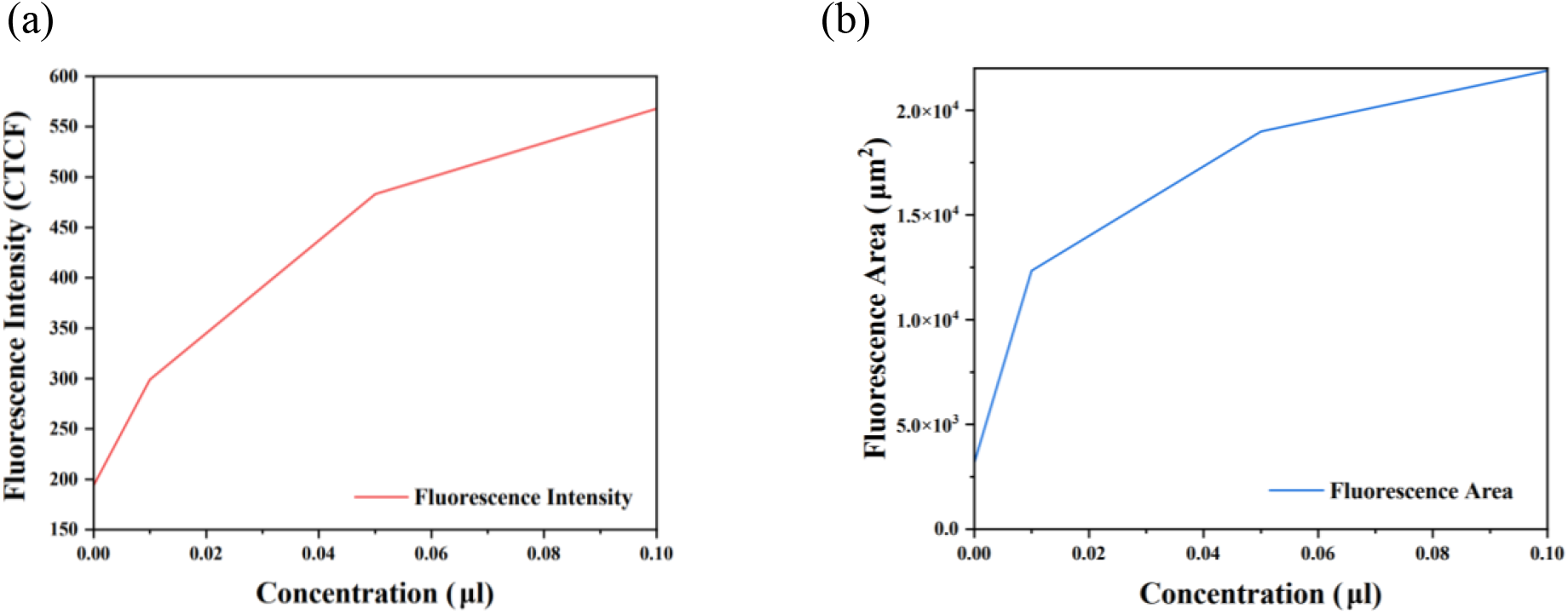
(a) CTCF graphed alginate beads versus quantum dot concentration, showing a logarithmic relationship. (b) Fluorescence area graphed versus concentration, as well showing a logarithmic relationship.

Next, we investigate the changes in the rate of photosynthesis at different concentrations of the quantum dots through a bicarbonate indicator solution. This solution comprises a pH indicator and HCO ^-^_3_ ions. At higher levels of CO_2_, the number of bicarbonate ions in the solution increases, resulting in a decrease in pH and a corresponding color change from orange to yellow hue. In contrast, lower levels of CO_2_ reduce the number of bicarbonates and thus increase pH. The solution color changes to a purple or blue hue. The characterization starts with the preparation of PCR tubes with 100 µL of bicarbonate indicator and 25 µL of CO_2_. The tubes were then mixed for 30 s, causing the red-hued indicator to turn yellow. Afterward, five beads were added to the PCR tubes, and the tubes were subsequently sealed and placed under a cool white fluorescent light for 2 hr. Every 30 min, the vials were mixed at low speed for 30 s and then photographed. As shown in **Figure 5**, the rate of pH change in the solution is lowered with increasing quantum dot concentration. Although qualitative, this visual observation confirms a negative correlation between quantum dot concentration and the rate of algae photosynthesis.

**Figure 5.**
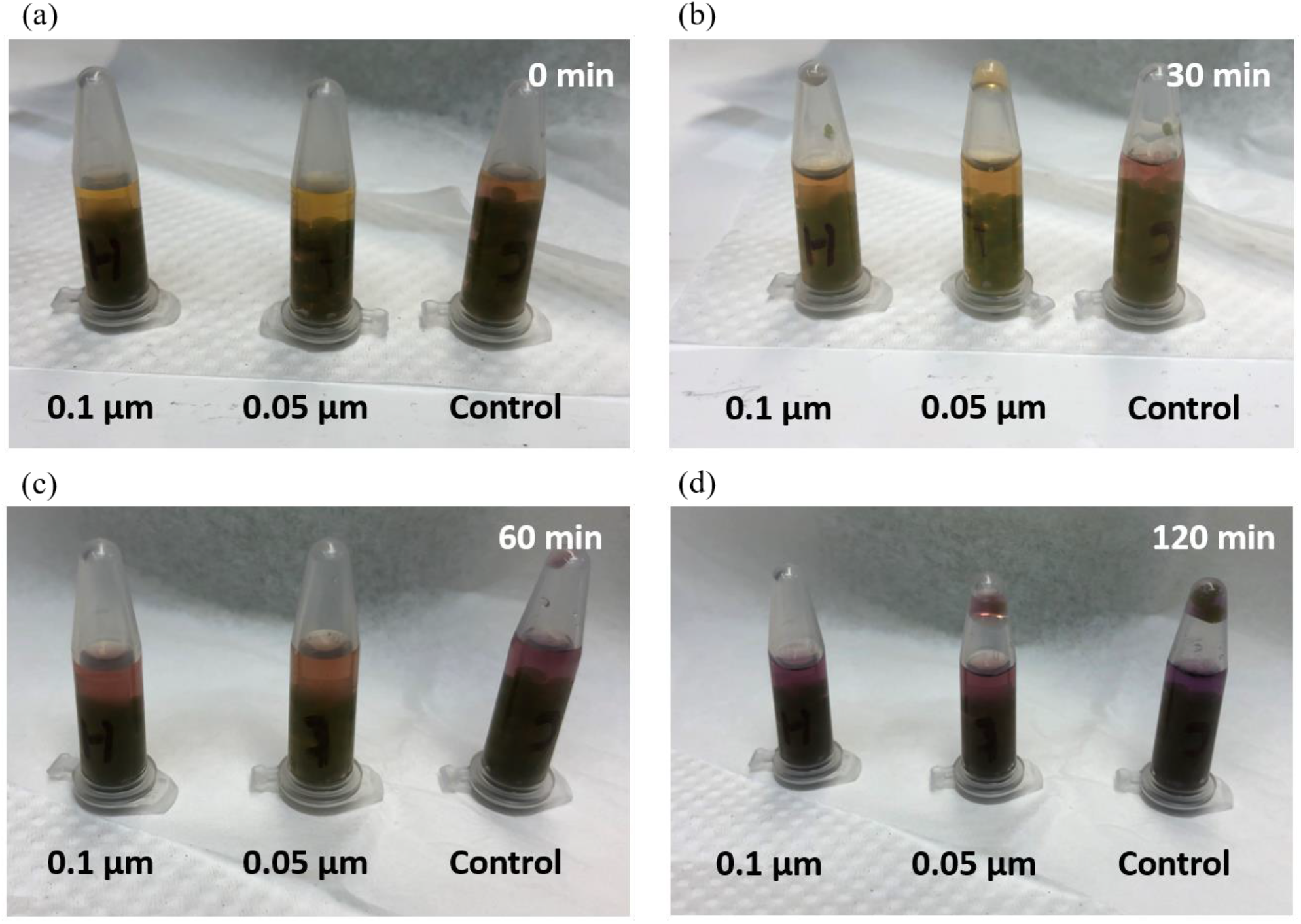
Qualitative analysis of the rate of photosynthesis using bicarbonate indicator, for quantum dots with a concentration ranging from 0 to 0.1 µM and an incubation time of (a) 0 min. (b) 30 min. (c) 60 min. (d) 120 min. The color of the solution indicates a change in CO_2_ concentration. The decrease in the CO_2_ level changes the solution from yellow to purple.

A Shimadzu gas chromatography-mass spectrometry was used to quantify the concentration of CO_2_ and O_2_ in the microalgae beads solution. Thirty-six chromatography vials were used to measure the concentrations every 24 min for 2 hr. Five beads with different concentrations of the quantum dots (0 uM, 0.01 uM, 0.05 uM, 0.1 uM) were added to the chromatography vials, and each vial was filled with 200 µL of water.Two hundred microliters of CO2 were added to the first set of vials at time 0, after which the vials were sealed and placed under a fluorescent lamp. At 24 min, 200 µL of CO_2_ were added to the second set of vials, and then they were placed under a fluorescent lamp. This process was repeated for all sets of vials until the last set produced at 120 min. The vials of the last set were filled with 200 µL of water but not placed under the fluorescent light. After 120 min, all vials were moved away from light and then taken to measure in the chromatography. The measurement of the time 0 set represents the final concentrations of gasses, while the set produced at 120 min gives the initial concentrations of gasses. As shown in **Figure 6a**, the CO_2_ concentration present in the sample over time is positively correlated with the concentration of quantum dots. On the other hand, the percent oxygen concentration present in the samples over time is negatively correlated with the concentration of quantum dots (**Figure 6b**). These quantitative data confirm a decrease in the rate of photosynthesis as the concentration of quantum dots increases.

**Figure 6.**
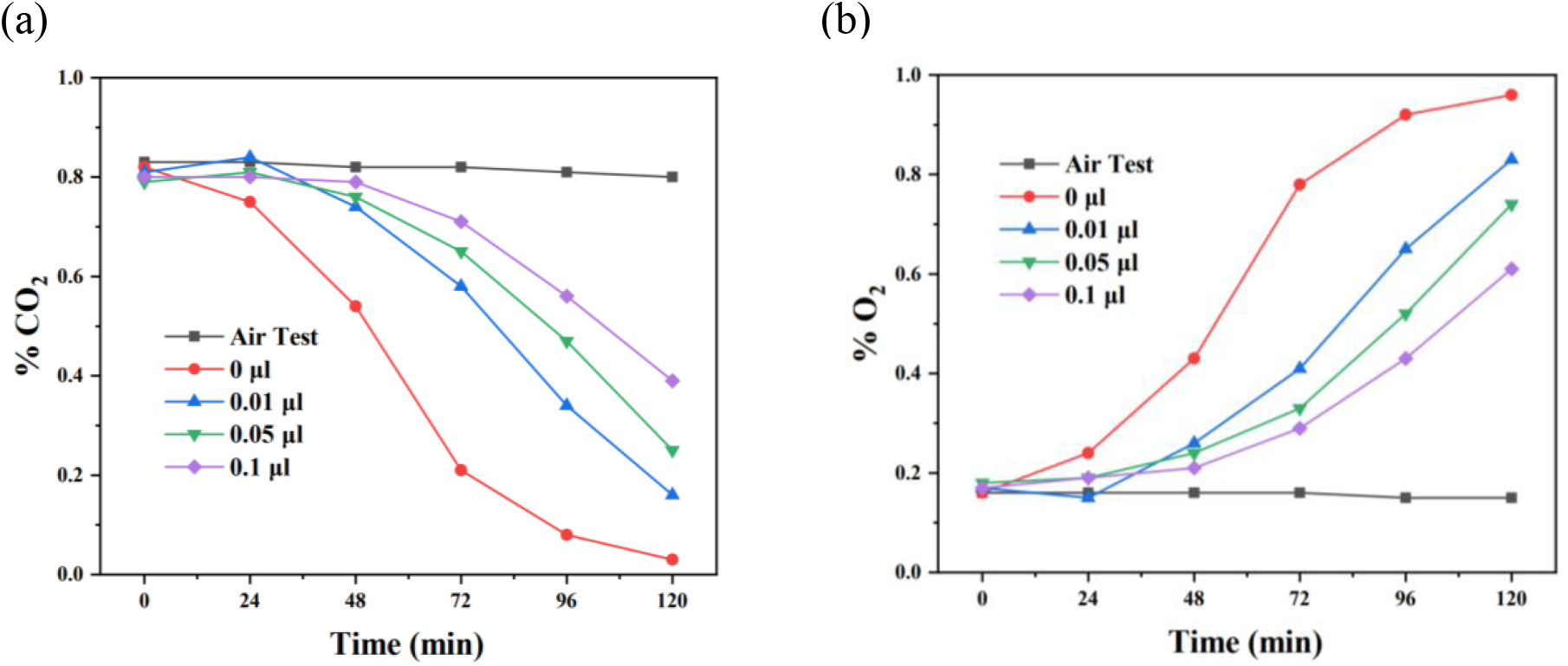
(a) CO_2_ and (b) O_2_ level versus time of the alginate beads solution with different concentrations of the quantum dots.

The scale-up of microalgae cultivation with photosynthesis requires living systems based on microalgae cells and polymer matrix to be created with high spatial cell densities^8^. However, the influence of engineering nanomaterials and debris on these living systems has not been explored^15,20^. Despite that alginate and other polymer building blocks can immobilize the cells and impede the quantum dots uptake^21^, we have found that the photosynthesis of the algae beads remains significantly affected, especially within the first 48 min after the incubation. Previous studies have found a correlation between the introduction of nanostructures to aquaponics plants and the inhibition of the cell wall functions^22,23^. The quantum dots could have similar adverse effects by inhibiting the vital movement of resources needed for photosynthetic activity in the algae beads, thus indirectly causing the plants’ rate of photosynthesis to decrease. Another possible cause is that the quantum dots directly interfere with the cells’ light absorption in the matrix^24^. This mechanism would decrease photosynthesis, but not cellular respiration, unlike the inhibition of the cell walls.

We found that the photosynthetic rate increases over time for the algae beads incubated with the quantum dots, regardless of the quantum dot concentration. For example, at 72 min, the O_2_ level is only 0.3% for the 0.1 µL sample, compared to 0.8% without quantum dots. On the other hand, at 120 min, the difference between the two samples decreases to ∼0.3%. As the pore size on the cell wall is 5-20 nm, the quantum dots can enter the cell wall through endocytosis to damage the cell structure, thus lowering the photosynthetic efficiency. However, the ubiquitous rate increase for all the samples indicates that the cells can adapt to the hazardous environment and still perform photosynthesis by either exocytosis or quantum dot digestion^25,26^.

## Acknowledgements

This material is based upon work supported by the National Science Foundation under Grant 2034855 to Xin Yong and 2035623 to Ke Du.

The authors would like to thank Xian Boles for the schematic design.

## References

(1) Tang, J.; Liang, Q.; Li, C.; Huang, X.; Xian, X.; Li, J.; Shang, Z.; Pang, C.; Liu, Y.; Zhang, R. Application of Marine Algae in Water Pollution Control. IOP Conf. Ser.: Earth Environ. Sci. 2022, 966 (1), 012001. https://doi.org/10.1088/1755-1315/966/1/012001.

(2) Adeniyi, O. M.; Azimov, U.; Burluka, A. Algae Biofuel: Current Status and Future Applications. Renewable and Sustainable Energy Reviews 2018, 90, 316–335. https://doi.org/10.1016/j.rser.2018.03.067.

(3) Bito, T.; Okumura, E.; Fujishima, M.; Watanabe, F. Potential of Chlorella as a Dietary Supplement to Promote Human Health. Nutrients 2020, 12 (9), 2524. https://doi.org/10.3390/nu12092524.

(4) Chapman, R. L. Algae: The World’s Most Important “Plants”—an Introduction. Mitig Adapt Strateg Glob Change 2013, 18 (1), 5–12. https://doi.org/10.1007/s11027-010-9255-9.

(5) Elliff, C. I.; Silva, I. R. Coral Reefs as the First Line of Defense: Shoreline Protection in Face of Climate Change. Marine Environmental Research 2017, 127, 148–154. https://doi.org/10.1016/j.marenvres.2017.03.007.

(6) Hoegh-Guldberg, O. Climate Change, Coral Bleaching and the Future of the World’s Coral Reefs. Mar. Freshwater Res. 1999, 50 (8), 839–866. https://doi.org/10.1071/mf99078.

(7) Buerger, P.; Alvarez-Roa, C.; Coppin, C. W.; Pearce, S. L.; Chakravarti, L. J.; Oakeshott, J. G.; Edwards, O. R.; van Oppen, M. J. H. Heat-Evolved Microalgal Symbionts Increase Coral Bleaching Tolerance. Science Advances 2020, 6 (20), eaba2498. https://doi.org/10.1126/sciadv.aba2498.

(8) Wangpraseurt, D.; You, S.; Azam, F.; Jacucci, G.; Gaidarenko, O.; Hildebrand, M.; Kühl, M.; Smith, A. G.; Davey, M. P.; Smith, A.; Deheyn, D. D.; Chen, S.; Vignolini, S. Bionic 3D Printed Corals. Nat Commun 2020, 11 (1), 1748. https://doi.org/10.1038/s41467-020-15486-4.

(9) Riera, E.; Lamy, D.; Goulard, C.; Francour, P.; Hubas, C. Biofilm Monitoring as a Tool to Assess the Efficiency of Artificial Reefs as Substrates: Toward 3D Printed Reefs. Ecological Engineering 2018, 120, 230–237. https://doi.org/10.1016/j.ecoleng.2018.06.005.

(10) Balasubramanian, S.; Yu, K.; Meyer, A. S.; Karana, E.; Aubin-Tam, M.-E. Bioprinting of Regenerative Photosynthetic Living Materials. Advanced Functional Materials 2021, 31 (31), 2011162. https://doi.org/10.1002/adfm.202011162.

(11) Marangoni, L. F. B.; Beraud, E.; Ferrier-Pagès, C. Polystyrene Nanoplastics Impair the Photosynthetic Capacities of Symbiodiniaceae and Promote Coral Bleaching. Science of The Total Environment 2022, 815, 152136. https://doi.org/10.1016/j.scitotenv.2021.152136.

(12) Ripken, C.; Khalturin, K.; Shoguchi, E. Response of Coral Reef Dinoflagellates to Nanoplastics under Experimental Conditions Suggests Downregulation of Cellular Metabolism. Microorganisms 2020, 8 (11), 1759. https://doi.org/10.3390/microorganisms8111759.

(13) Lin, S.; Bhattacharya, P.; Rajapakse, N. C.; Brune, D. E.; Ke, P. C. Effects of Quantum Dots Adsorption on Algal Photosynthesis. J. Phys. Chem. C 2009, 113 (25), 10962–10966. https://doi.org/10.1021/jp904343s.

(14) Yang, W.; Gao, P.; Li, H.; Huang, J.; Zhang, Y.; Ding, H.; Zhang, W. Mechanism of the Inhibition and Detoxification Effects of the Interaction between Nanoplastics and Microalgae Chlorella Pyrenoidosa. Science of The Total Environment 2021, 783, 146919. https://doi.org/10.1016/j.scitotenv.2021.146919.

(15) Khoshnamvand, M.; Hanachi, P.; Ashtiani, S.; Walker, T. R. Toxic Effects of Polystyrene Nanoplastics on Microalgae Chlorella Vulgaris: Changes in Biomass, Photosynthetic Pigments and Morphology. Chemosphere 2021, 280, 130725. https://doi.org/10.1016/j.chemosphere.2021.130725.

(16) Li, X.; Sun, H.; Mao, X.; Lao, Y.; Chen, F. Enhanced Photosynthesis of Carotenoids in Microalgae Driven by Light-Harvesting Gold Nanoparticles. ACS Sustainable Chem. Eng. 2020, 8 (20), 7600–7608. https://doi.org/10.1021/acssuschemeng.0c00315.

(17) Ward’s World Activity Guides - How to Make Your Own Algae Beads. https://wardsworld.wardsci.com/i/1144195-how-to-make-your-own-algae-beads/0? (accessed 2022-05-31).

(18) Allard, B.; Templier, J. Comparison of Neutral Lipid Profile of Various Trilaminar Outer Cell Wall (TLS)-Containing Microalgae with Emphasis on Algaenan Occurrence. Phytochemistry 2000, 54 (4), 369–380. https://doi.org/10.1016/S0031-9422(00)00135-7.

(19) Chen, P.; Powell, B. A.; Mortimer, M.; Ke, P. C. Adaptive Interactions between Zinc Oxide Nanoparticles and Chlorella Sp. Environ. Sci. Technol. 2012, 46 (21), 12178–12185. https://doi.org/10.1021/es303303g.

(20) Reichelt, S.; Gorokhova, E. Micro- and Nanoplastic Exposure Effects in Microalgae: A Meta-Analysis of Standard Growth Inhibition Tests. Frontiers in Environmental Science 2020, 8.

(21) Li, X.; Zhang, Z.; Xie, Q.; Wu, D. Effect of Algae on Phosphorus Immobilization by Lanthanum-Modified Zeolite. Environmental Pollution 2021, 276, 116713. https://doi.org/10.1016/j.envpol.2021.116713.

(22) Ji, J.; Long, Z.; Lin, D. Toxicity of Oxide Nanoparticles to the Green Algae Chlorella Sp. Chemical Engineering Journal 2011, 170 (2), 525–530. https://doi.org/10.1016/j.cej.2010.11.026.

(23) Yan, A.; Chen, Z. Impacts of Silver Nanoparticles on Plants: A Focus on the Phytotoxicity and Underlying Mechanism. International Journal of Molecular Sciences 2019, 20 (5), 1003. https://doi.org/10.3390/ijms20051003.

(24) Fan, G.; Chen, Z.; Yan, Z.; Du, B.; Pang, H.; Tang, D.; Luo, J.; Lin, J. Efficient Integration of Plasmonic Ag/AgCl with Perovskite-Type LaFeO3: Enhanced Visible-Light Photocatalytic Activity for Removal of Harmful Algae. Journal of Hazardous Materials 2021, 409, 125018. https://doi.org/10.1016/j.jhazmat.2020.125018.

(25) Chen, F.; Xiao, Z.; Yue, L.; Wang, J.; Feng, Y.; Zhu, X.; Wang, Z.; Xing, B. Algae Response to Engineered Nanoparticles: Current Understanding, Mechanisms and Implications. Environmental Science: Nano 2019, 6 (4), 1026–1042. https://doi.org/10.1039/C8EN01368C.

(26) Pulido-Reyes, G.; Marie Briffa, S.; Hurtado-Gallego, J.; Yudina, T.; Leganés, F.; Puntes, V.; Valsami-Jones, E.; Rosal, R.; Fernández-Piñas, F. Internalization and Toxicological Mechanisms of Uncoated and PVP-Coated Cerium Oxide Nanoparticles in the Freshwater Alga Chlamydomonas Reinhardtii. Environmental Science: Nano 2019, 6 (6), 1959–1972. https://doi.org/10.1039/C9EN00363K.

